# Predicting plasmodesmata-mediated interface permeability and intercellular diffusion

**DOI:** 10.1101/2024.08.06.606906

**Authors:** Johannes Liesche, Jiazhou Li, Helle Jakobe Martens, Chen Gao, Angeline Christina Subroto, Alexander Schulz, Eva Deinum

**Affiliations:** Institute of Biology, University of Graz, 8010 Graz, Austria; College of Life Sciences, Northwest A&F University, 712100 Yangling, China; Department of Geosciences and Natural Resource Management, University of Copenhagen, 1958 Frederiksberg, Denmark; Department of Plant and Environmental Sciences, University of Copenhagen, 1781 Frederiksberg, Denmark; Mathematical and Statistical Methods (Biometris), Plant Science Group, Wageningen University, 6708PB Wageningen, The Netherlands

**Author notes:** Corresponding author: Prof. Dr. Johannes Liesche; Institute of Biology, University of Graz, Schubertstraße 51A, 8010 Graz, Austria.

**Keywords:** Intercellular transport, electron microscopy, theoretical modeling, uncaging, diffusion, glucosinolate, Ca^2+^ signaling, MYC2

## Abstract

Intercellular communication is essential for plant development and responses to biotic and abiotic stress. A key pathway is diffusive exchange of signal molecules and nutrients via plasmodesmata. These cell wall channels connect the cytoplasms of most cells in land plants. Their small size, with a typical diameter of about 50 nm, and complex structure have hindered the quantification plasmodesmata-mediated intercellular diffusion. This measure is essential for disentangling the contributions of diffusive and membrane transporter-mediated movement of molecules that, together, define cell interactions within and across tissues. We compared the two most promising methods to measure plasmodesmata-mediated interface permeability, live-cell microscopy with fluorescent tracer molecules and transmission electron microscopy-based mathematical modeling, to evaluate the potential for obtaining absolute quantitative values. We applied both methods to 29 cell-cell interfaces from nine angiosperm species and found a stronger association between the modelled and experimentally determined interface permeabilities than between the experimentally-determined permeability and any single structural parameter. By feeding the values into a simulation of an artificial Arabidopsis leaf, we illustrate how interface permeabilities can help to predict diffusion patterns of defense-related molecules, such as glucosinolates and transcription factors.

## Introduction

Plasmodesmata (PD) are cell wall channels that provide a degree of cytosolic continuity between most plant cells. PD structure and abundance influence which molecules can move intercellularly and at what rate. The PD-mediated transport capacity is critical for processes such as organ development (Otero et al. 2016; Godel-Jedrychowska et al. 2021), pathogen defense (Cheval and Faulkner 2018, Li et al. 2021), environmental acclimation (Bilska and Sowinski 2010; Cui and Lee 2016; Gao et al. 2020), and carbon allocation (Comtet et al. 2017; Liesche 2017; Miras et al. 2022). Accordingly, manipulation of PD has been proposed as a strategy to optimize biotic stress resistance (Liu et al. 2021) and crop productivity (Sun et al. 2019; Amari et al. 2021).

The main route for molecular movement through PD is the cytoplasmic sleeve, the space between the plasma membrane and the ER membrane that forms a central tube called desmotubule. The width of the cytoplasmic sleeve poses a size-exclusion limit that typically prevents medium- and large-sized proteins from passing through PD (Tee and Faulkner 2024). Cell differentiation programs or specific molecular or physical cues can cause the cytoplasmic sleeve to widen, narrow or even close completely (Benitez-Alfonso et al. 2013). In most cases, where this has been analyzed, the variation in cytoplasmic sleeve width, especially at the neck regions of PD, is due to callose deposition or breakdown in the adjacent cell wall (Amsbury et al. 2018). Some indications of a callose-independent regulation have been presented (Kitagawa et al. 2019). The variation of cytoplasmic sleeve width does not only change the size-exclusion limit, but also affects the rate at which molecules of permitted size pass through PD (Deinum et al. 2019; Ostermeyer et al. 2022). Determining this rate is essential for a quantitative understanding of inter-cellular transport (Sowinski et al. 2008; Tee and Faulkner 2024). However, absolute quantification of PD transport is challenging. Single PD are too small to be observed under a conventional light microscope and their position within the cell wall makes them inaccessible to live-cell super-resolution microscopy (Liesche et al. 2013). Therefore, instead of the rate of molecular movement through single PD, cell wall permeability mediated by all PD in a cell-cell interface has been established as a variable for assessing intercellular movement (Rutschow et al. 2011). We will refer to this PD-mediated interface permeability as ‘interface permeability’ or ‘effective permeability’ (P_eff_) in the following to avoid confusion with the permeability of individual PD. Interface permeability P_eff_ can be used to calculate flux F, i. e. the number of molecules per unit surface area moving through PD from one side of the cell wall to the other side in a given time period, when the cytosolic concentration potential *Δc* on both ends of the PD is known, with *F* = *P*_*eff*_ ∗ Δ*c*.

Radioactively labeled molecules have been used to assess PD function (Pitman 1971; Kwiatkowska 1991; Schulz 1995) and could be used to measure interface permeability. However, working with radioactive substances has drawbacks, such as difficulty with targeted detection as well as issues of safety and regulation. Instead, fluorescent tracers have been applied for this purpose, especially fluorescein. Fluorescein has a similar size to simple sugars and plant hormones allowing for conclusions regarding their movement based on the measurement of fluorescein movement (Liesche and Schulz 2012; Gao et al. 2020). Quantifying P_eff_ using fluorescent tracers typically requires confocal microscopy and photobleaching or photoactivation protocols for sufficient temporal and spatial resolution (Liesche and Schulz 2012; Martens et al. 2019). While this approach offers a low level of invasiveness, the analysis is limited to cells and tissues that are accessible to confocal microscopy. In practice, such experiments have been almost exclusively conducted on epidermis cells (Rutschow et al. 2011; Gao et al. 2020).

The alternative to calculating interface permeability based on intercellular tracer movement is a calculation based on structural data, obtained from transmission electron microscopy (TEM). The measurement of PD abundance from a series of TEM sections has been used to estimate interface permeability (Botha and van Bel 1992; Liesche et al. 2019; Botha and Murugan 2021). Liesche et al. (2019) took PD structural parameters, such as neck diameter, into account for a comparative analysis of different cell-cell interfaces, but found that these parameters did not correlate with tracer-based measures of permeabilities. A recent biophysical modeling approach of molecular movement through PD enabled the prediction of interface permeability based on various structural features extracted from TEM images (Deinum et al. 2019). The predictions roughly matched interface permeability values calculated from fluorescence tracer approaches in the Arabidopsis root meristem (Deinum et al. 2019). The calculations have been compiled in the open-source software PDinsight (Deinum 2022).

It should be noted that both methods of calculating absolute values of interface permeability, from fluorescent tracer movement and from PD structure, have technical limitations that introduce an error. The magnitude of this error could not be conclusively assessed (Liesche and Schulz 2012; Deinum et al. 2019), casting some doubt on the absolute values resulting from both methods. For the tracer movement, the measurement of concentrations at the PD openings is limited by the temporal and spatial resolution of the microscope and assumptions of homogeneity in the data analysis (Liesche and Schulz 2012). Accordingly, tracer concentrations are typically averaged over several seconds and several µm, leading to variations of P values of the same interface of up to 50 % (Liesche and Schulz 2012). For the TEM-based calculations, there is a potential influence of structural elements that are not resolved on images, e.g. proteins lowering the diffusivity in the cytosolic sleeve (Liesche and Schulz 2013). Moreover, PD are generally assumed to be very sensitive to tissue disturbances (Oparka and Prior 1992; Park et al. 2019), raising the possibility of structural aberrations introduced by the highly invasive TEM sample preparation process (Peters et al. 2021).

With this study, we have two aims: i) to evaluate the relationship between interface permeabilities derived from TEM-based calculations and fluorescent tracer approaches; ii) to illustrate how interface permeabilities can help to predict diffusion patterns of various molecules, using the Arabidopsis leaf as example.

Supporting these aims, we quantify desmotubule structure on TEM images. The desmotubule is generally difficult to discern on two-dimensional TEM images and, accordingly, there has been no systematic analysis of its structural variability and its potential link to other PD parameters. This is despite desmotubule structure, especially its diameter, carrying a high significance for the calculation of PD permeability (Deinum et al. 2019; Park et al. 2019; Peters et al. 2021). Our data on desmotubule structure informs the following calculations of interface permeability in PDinsight.

## Results

### Relationship of desmotubule diameter to PD structure

The desmotubule is not visible on most images produced using standard TEM methods. Of the 523 PD imaged in this study, only 29 showed a distinct desmotubule (Fig. 1). One reason is that they typically cover longitudinal views of PD (Fig. 1A,B). For some interfaces, we produced transverse sections to verify desmotubule radii (Fig. 1C). Our data was supplemented with literature data to represent a total of 69 PD from various cell-cell interfaces in 20 plant species (Table S1). Desmotubule radius data was used to test for correlations with other PD-related structural parameters, namely neck radius, neck length, max radius (typically at the central part of a PD) and length.

**Fig. 1.**
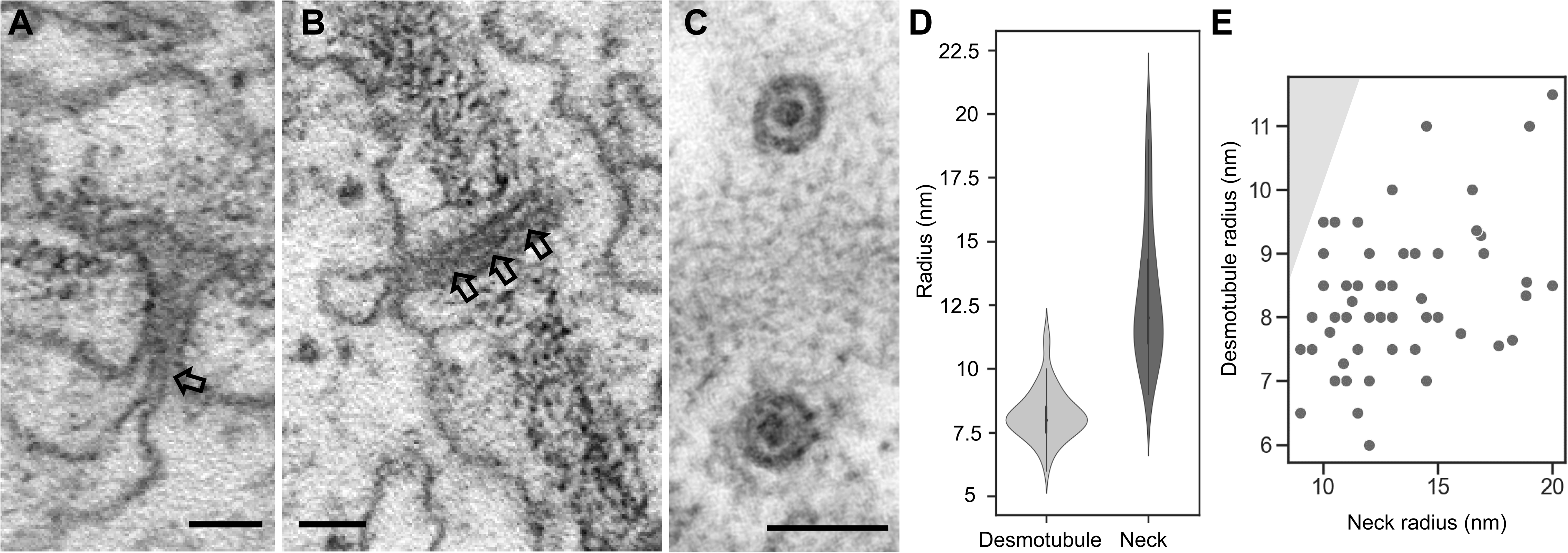
The structure of primary plasmodesmata (PD) desmotubules. **A, B** Images of PD from Arabidopsis (**A**) and *Cucurbita maxima* (**B**) illustrate the appearance of desmotubules (arrows) on conventional transmission electron micrographs. Desmotubules can mostly be recognized at the neck regions, where they connect to the ER (**A**). In some cases, the desmotubule is visible along the whole length of the PD (**B**). **C** Transverse sections provide a clearer view of plasma membrane (outer ring), desmotubule (inner circle) and the cytosolic sleeve (space between plasma membrane and desmotubule) of PDs from *Pisum sativum*. Scale bars 50 nm. **D** Distribution of radii of desmotubules and PD necks. **E** Desmotubule radii plotted against PD neck radii. Grey shading indicates the biologically nonsensical chart area where desmotubule radius would be larger than neck radius.

The average desmotubule radius was 8.25 nm. Notable is the small variance of desmotubule radii. While neck radii displayed variance of about 60 %, desmotubule radii of the same PDs vary only by about 12 % (Fig. 1D, Table S1). Significant correlations were detected between desmotubule radius and PD neck radius and max radius (Table 1). The strongest correlation was found between desmotubule radius and PD neck radius with a correlation coefficient of 0.42 (Table 1, Fig. 1E).

**Table 1.**
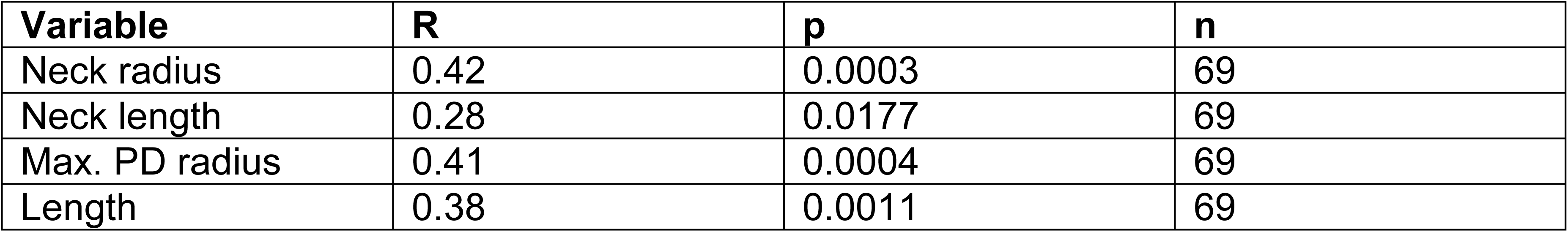
Pearson correlation statistics of the relationship between desmotubule radius and plasmodesmata structural variables. The analysis includes 40 plasmodesmata from literature sources. The Bonferroni-threshold for significance with α = 0.05 is p < 0.0007. Underlying data is provided in Table S1.

| Variable | R | p | n |
| --- | --- | --- | --- |
| Neck radius | 0.42 | 0.0003 | 69 |
| Neck length | 0.28 | 0.0177 | 69 |
| Max. PD radius | 0.41 | 0.0004 | 69 |
| Length | 0.38 | 0.0011 | 69 |

### Comparison of interface permeabilities from modeling and photobleaching experiments

Our collection of TEM images contained 259 PDs at cell-cell interfaces that have been previously probed by fluorescence loss in photobleaching (FLIP) experiments (Liesche et al. 2019). FLIP experiments measure the ability of the small tracer carboxyfluorescein to move between cells by quantifying fluorescence reduction in the cells adjacent to the cell in which the fluorescent tracer is bleached. Due to the low time resolution of these experiments, we did not calculate absolute interface permeability values. Instead, we used the degree of fluorescence reduction normalized to interface area (FLIP value) as indicator of cell coupling. For the TEM data, relevant parameters were quantified (Table S2) and used to calculate permeability with PDinsight. Since desmotubule radius could not be determined for most PD, the average desmotubule radius of 8.25 nm determined above, was assumed. Small variation in desmotubule radii (7 nm or 9.5 nm) had a negligible effect on Peff calculations except for the specialized interface between bundle sheath cells and intermediary cells in cucurbits that features very narrow PDs (Fig. S1). The combined data represents eight plant species, each with two interfaces: i) mesophyll cell to bundle sheath cell and ii) bundle sheath cell to vascular parenchyma cell (Table 2). For *Primula vulgaris*, data was only available for ii).

**Table 2.** Data on plasmodesmata-related parameters from electron microscopy analysis in relation to modelled interface permeability (Model P_eff_) and FLIP value. FLIP experiments were carried out on living cells, making the resulting FLIP value an indicator of *in vivo* interface permeability. MC – mesophyll cell, BSC – bundle sheath cell, VP – vascular parenchyma cell. Underlying data of the TEM analysis is provided in Table S2, full results of P_eff_ modeling in Table S3.

| Species | Interface | Length (nm) | Max radius (nm) | Neck radius (nm) | Neck length (nm) | Density (PD $\mu\text{m}^{-2}$) | $N_{\text{TEM}}$ | Model $P_{\text{eff}}$ ($\mu\text{m s}^{-1}$) | FLIP value ($\mu\text{m s}^{-1}$) | $N_{\text{FLIP}}$ |
| --- | --- | --- | --- | --- | --- | --- | --- | --- | --- | --- |
| *Aesculus hippocastaneum* | MC-BSC | 202.33 | 33.43 | 27.00 | 27.63 | 4.75 | 15 | 8.05 | 44.6 | 3 |
|  | BSC-VP | 250.52 | 30.45 | 19.57 | 31 | 4.90 | 21 | 5.32 | 46.4 | 3 |
| *Betula pubescens* | MC-BSC | 337.15 | 31.15 | 22.91 | 35.35 | 2.54 | 20 | 2.23 | 28.2 | 12 |
|  | BSC-VP | 258.8 | 21.36 | 18.53 | 35.93 | 2.79 | 15 | 1.75 | 41.6 | 12 |
| *Malus domestica* | MC-BSC | 308.69 | 30.53 | 22.67 | 56 | 2.49 | 13 | 2.14 | 29.8 | 10 |
|  | BSC-VP | 293.05 | 22.075 | 19.35 | 35.35 | 2.1 | 20 | 1.07 | 39.5 | 10 |
| *Populus canescens* | MC-BSC | 392.12 | 27.22 | 17.46 | 59.22 | 7.87 | 22 | 3.19 | 31.7 | 10 |
|  | BSC-VP | 336.55 | 21.35 | 17.25 | 48.55 | 4.33 | 20 | 1.35 | 39.5 | 10 |
| *Nicotiana tabacum* | MC-BSC | 180.6 | 19 | 13.43 | 28.92 | 7.19 | 20 | 2.37 | 34.65 | 13 |
|  | BSC-VP | 175.41 | 19.41 | 14.51 | 20.75 | 2.35 | 12 | 1.10 | 18.05 | 13 |
| *Primula vulgaris* | BSC-VP | 311.45 | 23.32 | 17.31 | 30.90 | 1.51 | 11 | 0.72 | 14.7 | 5 |
| *Vicia faba* | MC-BSC | 206.73 | 20.7 | 14.88 | 34.8 | 3.72 | 15 | 1.46 | 31.5 | 10 |
|  | BSC-VP | 263.77 | 16.38 | 14.22 | 24.22 | 2.46 | 10 | 0.61 | 18 | 10 |
| *Cucurbita pepo* | MC-BSC | 182.6 | 15.86 | 12.26 | 24.13 | 8.02 | 15 | 1.50 | 31.1 | 15 |
|  | BSC-VP | 329.37 | 20.78 | 9.84 | 18.03 | 42.59 | 29 | 9.45 | 57.1 | 15 |

We observed no significant correlation of FLIP values with the individual structural parameters PD length, maximal radius, neck radius, neck length (p > 0.05) (Fig. 2A,B, Table 3). FLIP values correlated with PD density (Table 3), but this relationship was strongly influenced by a single datapoint corresponding to the specialized cucurbit interface (Fig. 2B). Without this point, the there is no significant correlation between the two parameters (R = 0.34; p = 0.23). In contrast, the effective permeability calculated in PDinsight based on TEM image-derived parameters correlated strongly with the FLIP value, with a correlation coefficient of 0.78 (p = 0.0006) (Fig. 2C, Table 3).

**Fig. 2.**
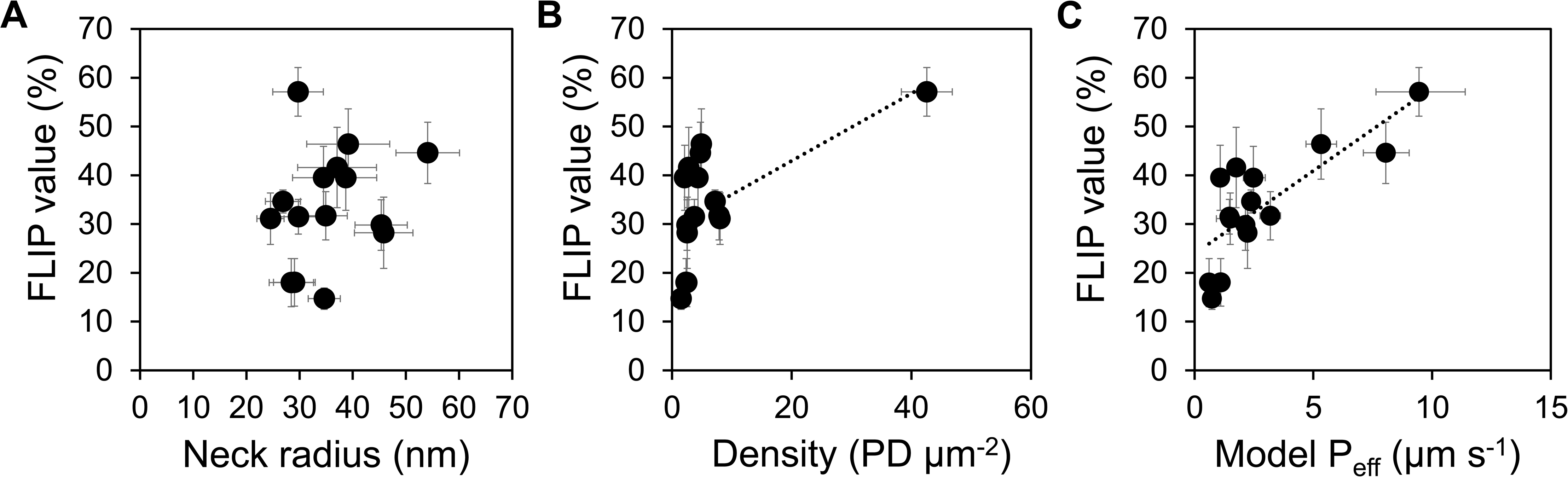
Relationship of fluorescence loss in photobleaching (FLIP) values to interface permeability calculated from electron microscopy images of plasmodesmata. **A** FLIP value plotted against plasmodesmata neck radius. **B** FLIP value plotted against plasmodesmata density. C FLIP value plotted against effective interface permeability calculated with PDinsight (Model P_eff_). Dotted line indicates significant correlation. The underlying data is provided in Table 2 and results of correlation analysis in Table 3.

**Table 3:** Pearson correlation statistics of the relationship between FLIP values and parameters determined from TEM images, as well as the effective interface permeability calculated in PDinsight (Model P_eff_). P-values < 0.05 are highlighted in bold. The analysis includes data for 15 interfaces (details in Table 2).

|  | R | p |
| --- | --- | --- |
| Density | 0.606 | **0.017** |
| Length | 0.103 | 0.715 |
| Max radius | 0.225 | 0.42 |
| Neck length | -0.039 | 0.89 |
| Neck radius | 0.232 | 0.41 |
| Model Peff | 0.781 | 0.0006 |

### Comparison of interface permeabilities from modeling and photoactivation experiments

Photoactivation experiments have been employed to determine absolute values of interface permeabilities at a wide range of cell types, especially in leaves (Liesche and Schulz 2012; Gao et al. 2020). The shorter time scales and higher contrast can increase reliability compared to photobleaching-based approaches (Liesche and Schulz 2012). Experiments were conducted on epidermis cells of Arabidopsis leaves at different developmental stages and Arabidopsis root cells (Fig. 3), *Cucurbita pepo* leaf mesophyll cells, as well as abaxial leaf epidermis cells of *Nicotiana tabacum* and *Populus x canescens* (Table 4). The results were supplemented with literature data for additional interfaces in Arabidopsis and *Cucurbita pepo* (Table 4). This included the interface between epidermis cells in the Arabidopsis *gsl8* mutant, which is deficient in callose synthesis and has, therefore, a higher PD permeability than wild-type plants (Gao et al. 2020). Like for FLIP values above, the actual P_eff_-values determined in photoactivation experiments were compared with data from TEM analysis, including modelled P_eff_ (Table 4, Fig. 3).

**Figure 3:**
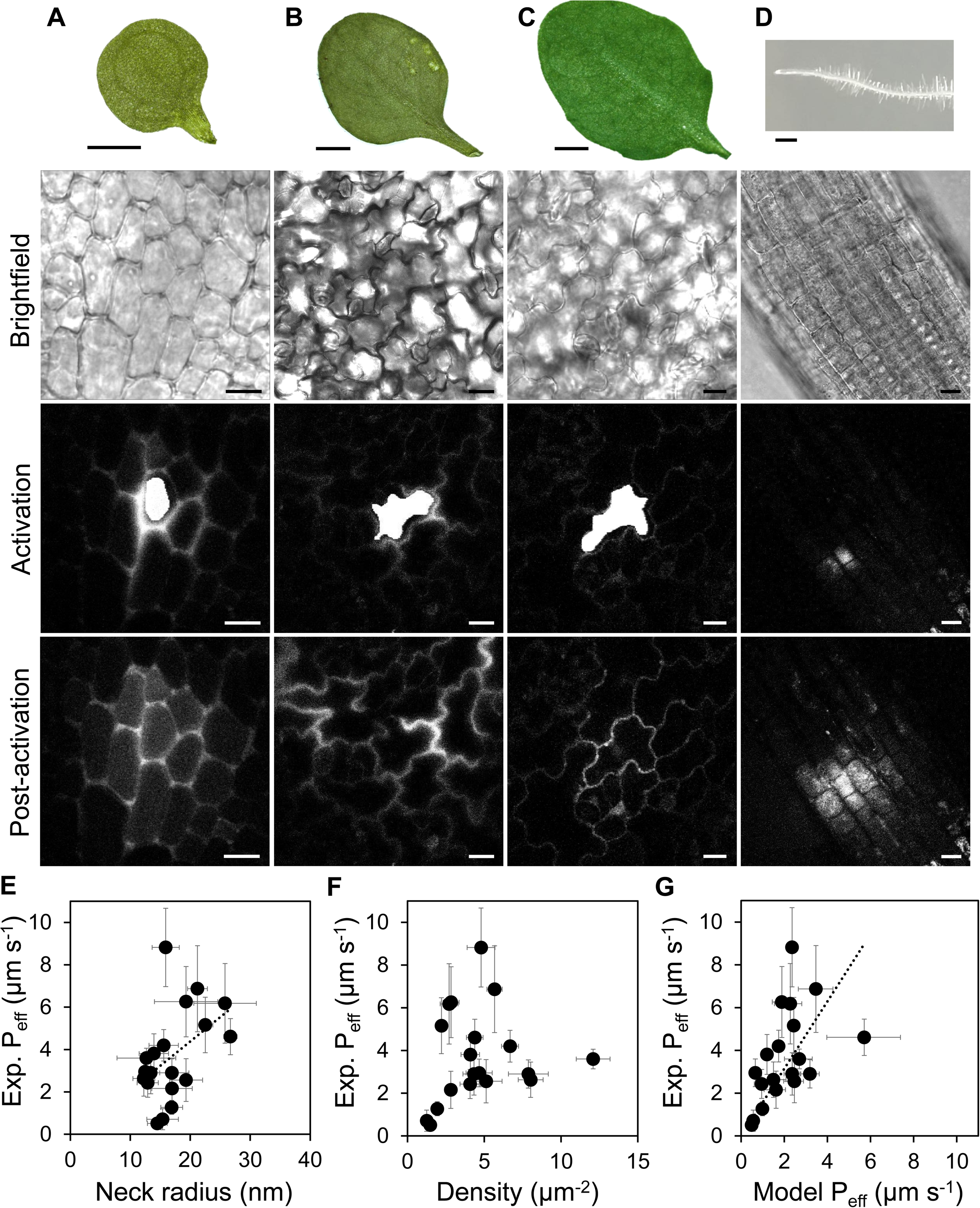
Measurement of interface permeability with live-cell photoactivation microscopy and relation to parameters derived from electron microscopy analysis. **A-D** Examples of photoactivation experiments conducted on abaxial epidermis cells of Arabidopsis leaves at young (**A**), transition (**B**) and mature (**C**) developmental stages and in Arabidopsis root cortex cells (**D**). The upper panels show the respective organ. Below is the bright field image that is used for orientation and the selection of a target cell. In this target cell, the fluorescent tracer Cage500 is activated, seen as white area on the fluorescence image in the panel below. In the lower panel, the first image after activation (Post-activation) is shown with visible spread of tracer from target to neighboring cells. The rate of tracer spread and fluorescence concentrations are used to calculate an effective interface permeability. Scale bars 1 mm (top panels), 10 µm (lower panels). **E-G** Correlation analysis between the interface permeability determined in photoactivation experiments and plasmodesmata neck radius (**E**), density (**F**) and interface permeability calculated in PDinsight based on electron microscopy data (**G**). Data includes measurements on wild-type and *gsl8* mutant plants. Error bars indicate standard deviation. Black dotted lines indicate significant Pearson correlation (P < 0.05). The corresponding data is provided in Table 4 and Table 5.

**Table 4.**
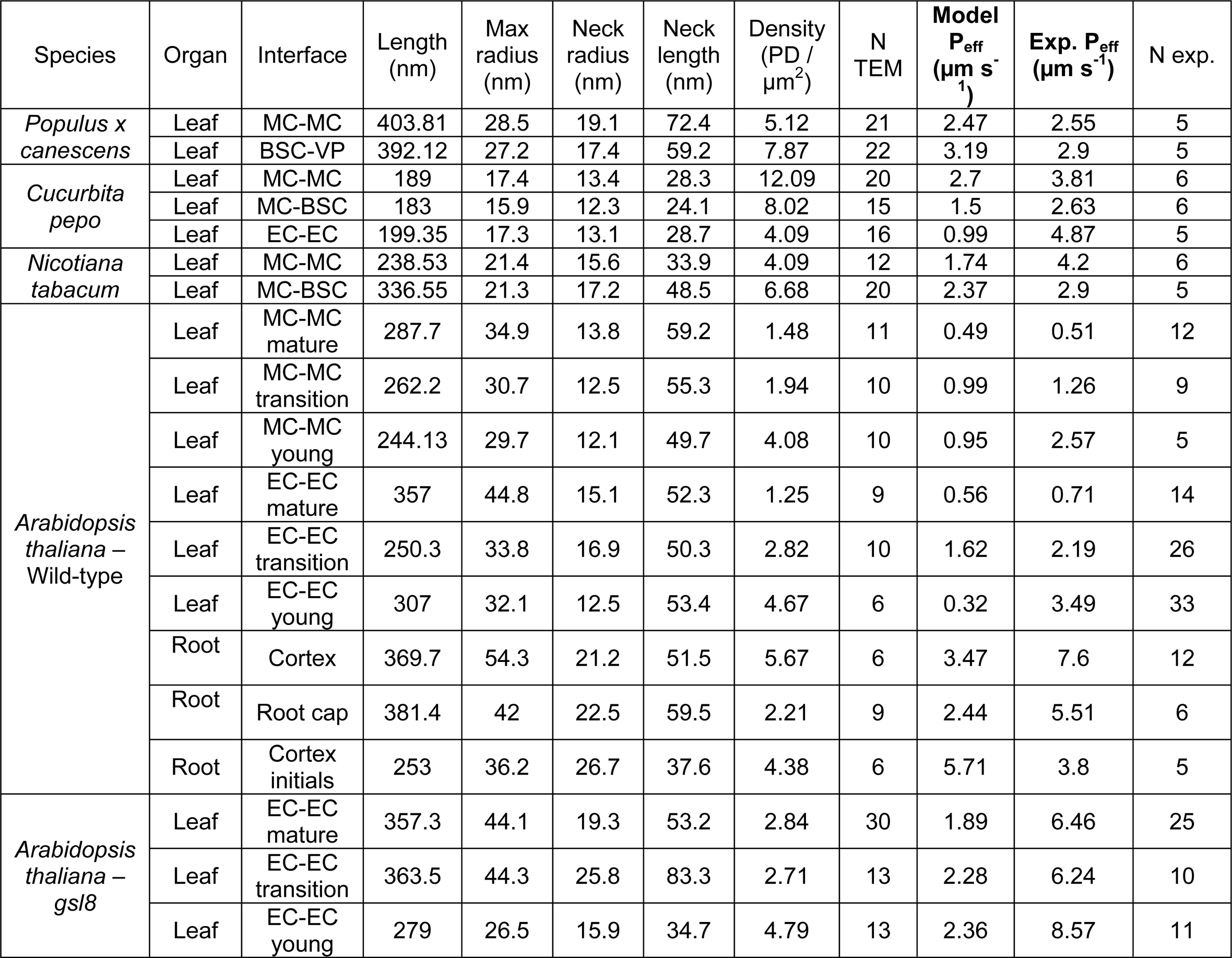
Data on plasmodesmata-related parameters from electron microscopy analysis in relation to modelled interface permeability (Model P_eff_) and interface permeability determined in photoactivation experiments (Exp. P_eff_). Underlying data of the TEM analysis is provided in Table S2, full results of P_eff_ modeling in Table S3.

**Table 5.**
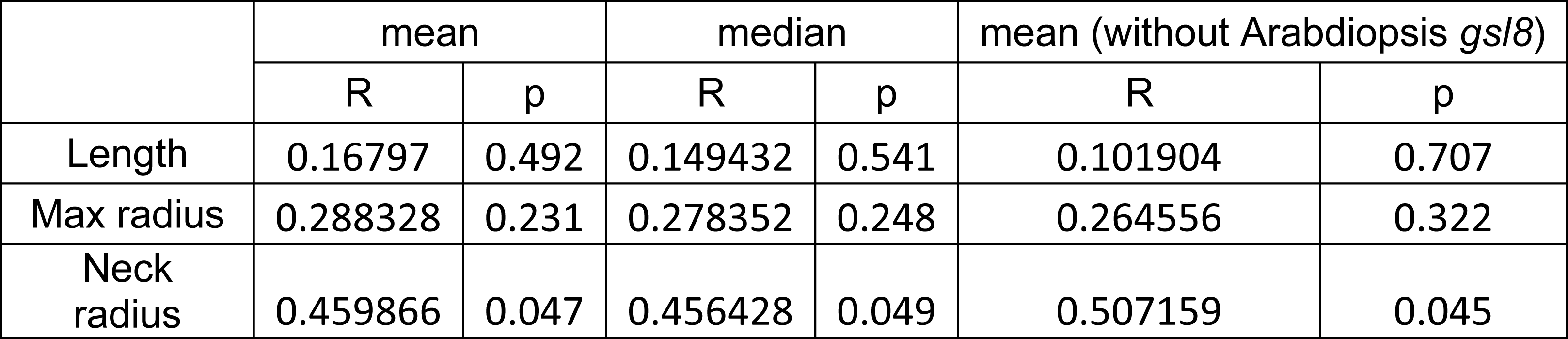
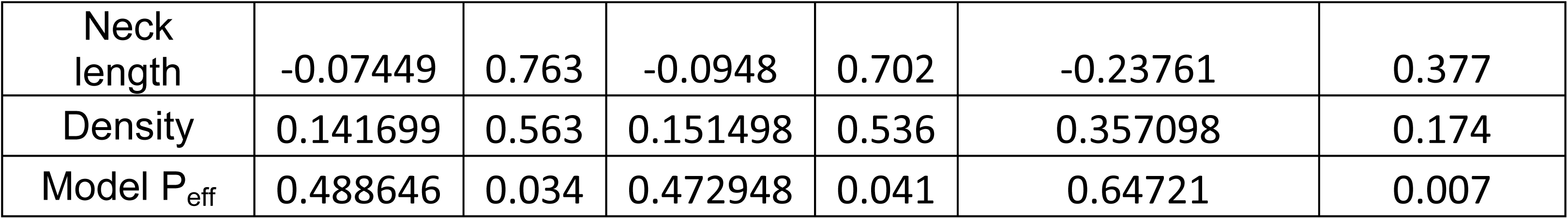
Pearson correlation statistics of the relationship between interface permeability determined by photoactivation experiments and parameters determined from TEM images, as well as the effective interface permeability calculated in PDinsight (Model P_eff_). P-values < 0.05 are highlighted in bold. The analysis includes data for 19 (all data) or 16 interfaces (without Arabidopsis *gsl8*) with details provided in Table 4.

The dataset represents interfaces with a wide range of PD structure and density as well as interface permeabilities (Table 4). Correlation analysis between experimentally determined interface permeability and the parameters determined from TEM images showed a correlation with neck diameter (Fig. 3E), but not PD density (Fig. 3F) (Table 5). Furthermore, the experimental P_eff_ correlated with the modelled P_eff_ (Fig. 3G), although with only slightly higher correlation coefficient than experimental P_eff_ and neck diameter (Table 5). Experimental P_eff_-values were about two times higher than modelled values (Fig. 3G). Correlation analysis with P_eff_ median values instead of mean values did not lead to different results (Table 5). However, excluding the three interfaces of the Arabidopsis *gsl8* mutant from the correlation analysis led to marked increases in the correlation coefficients of the associations between experimental Peff and PD density and between experimental P_eff_ and modelled P_eff_ (Table 5).

The comparison of interface permeabilities from photoactivation experiments and TEM data confirms the conclusion drawn from the comparison to FLIP results, that the modelled P_eff_ can be used to predict the *in situ* interface permeability, especially in wild-type plants.

### Interface permeabilities enable prediction of defense compound distributions

The interface permeability calculated from photoactivation experiments is specific for the activated tracer, in this case, a molecule with a hydrodynamic radius of about 0.6 nm (Belov et al. 2014). That means that the biological relevance of results is limited to similar molecules, for example sucrose (Liesche et al. 2019). In contrast, modeling the interface permeability based on PD structure and density data, as implemented in PDinsight, enables the consideration of molecules of various sizes (Deinum et al. 2019). We illustrated the potential of this approach by calculating interface permeabilities for molecules the size of three defense compounds with contrasting molecule sizes and simulating their movement in a simplified Arabidopsis leaf epidermis.

To obtain cell numbers and sizes, we conducted whole-mount staining with the cell wall-specific dye propidium iodide and fluorescence imaging of a transition-stage Arabidopsis leaf. Single images were stitched to yield a high-resolution view of the whole leaf (Fig. 4A-C). The average area of pavement cells was 2860 µm^2^ (Stdev = 488 µm, n = 300; Fig. 4B). Midrib cells were, on average, 112 µm long (stdev = 31.1 µm) and 20.5 µm wide (stdev = 4.3 µm) (Fig. 5C). For simplicity, only these two types of epidermis cells were considered in the model. In one leaf with an area of about 3.5 mm^2^, we counted about 940 pavement and 96 midrib cells (Fig. 4A). We designed the artificial leaf with a uniform arrangement of 50.6×50.6 µm^2^ pavement and similar width midrib cells with increasing aspect ratios towards the leaf base and petiole (Fig. 4D).

**Fig. 4:**
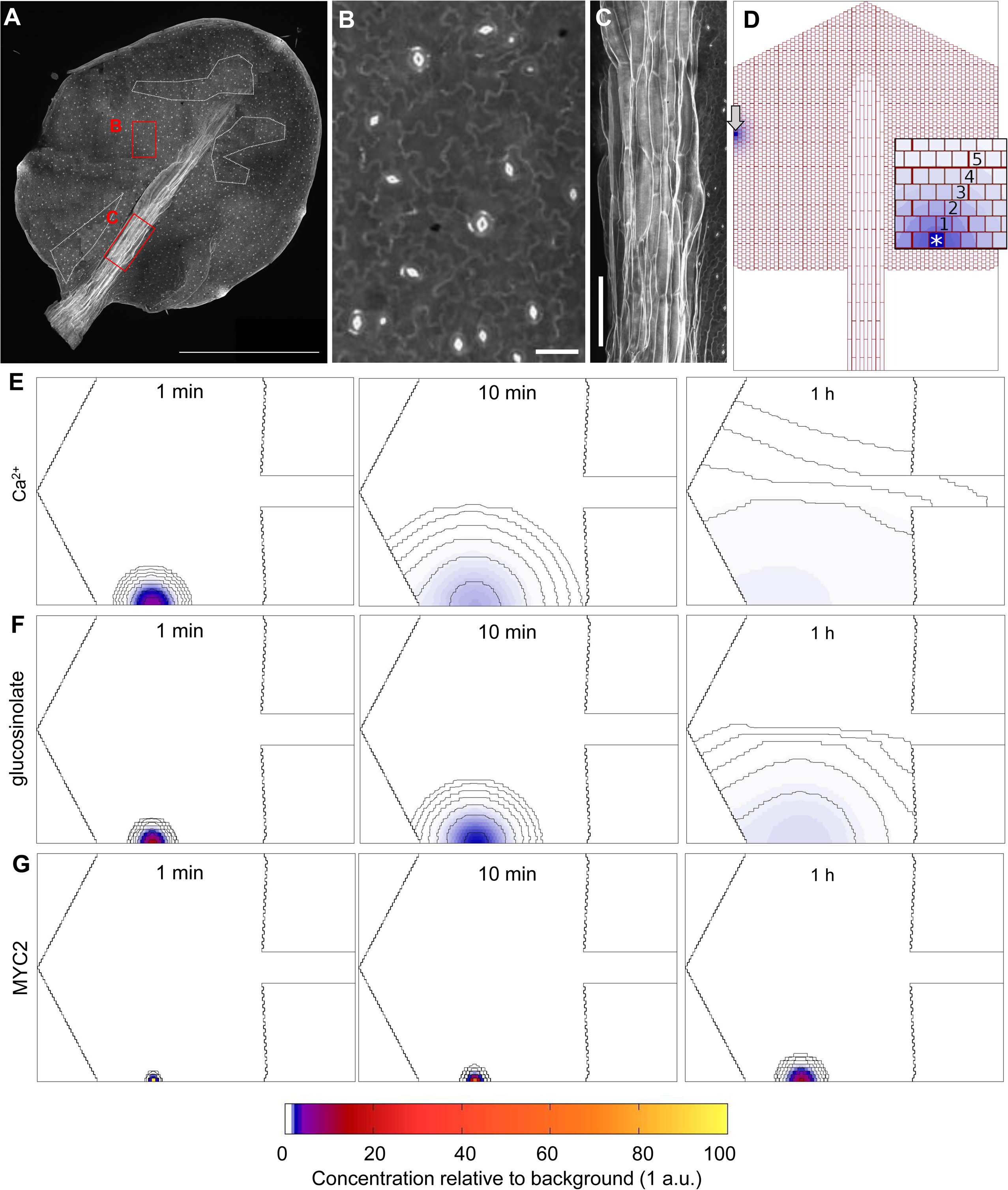
Theoretical modeling of the diffusion of defense-related compounds in Arabidopsis leaf epidermis. **A-C** Whole mount staining of Arabidopsis leaf. On the overview (**A**), mainly the walls of petiole and epidermis cells are visible, as well as the stomata (dots). The magnified views show the walls of pavement cells (**B**) and midrib epidermis (**C**). The overview image was stitched from 111 individual images. Outlined regions in **A** indicate areas that were not focused on original images and replaced with images from neighboring areas to provide a complete overview. Scale bars 1 mm (A), 20 µm (B), 100 µm (C) **D** Cell numbers and sizes from whole-mount staining analysis informed a simplified representation of the leaf epidermis. This was used for the following simulations of defense compound diffusion. **E-G** Simulations of diffusion of molecules with the size of Ca^2+^ (**E**), the glucosinolate 4-methylsulfinylpropyl (**F**) and MYC2 (**G**) in the stylized epidermis of an Arabidopsis leaf. Circular lines around the source cell are contour lines indicating relative increase above background levels (background concentration: 1 a.u.). Contours are spaced logarithmically, at 0.1%, 0.3%, 1%, 3% etc above background. Details of the model, anatomical assumptions and interface permeabilities are provided in the results section of the main text.

**Fig. 5.**
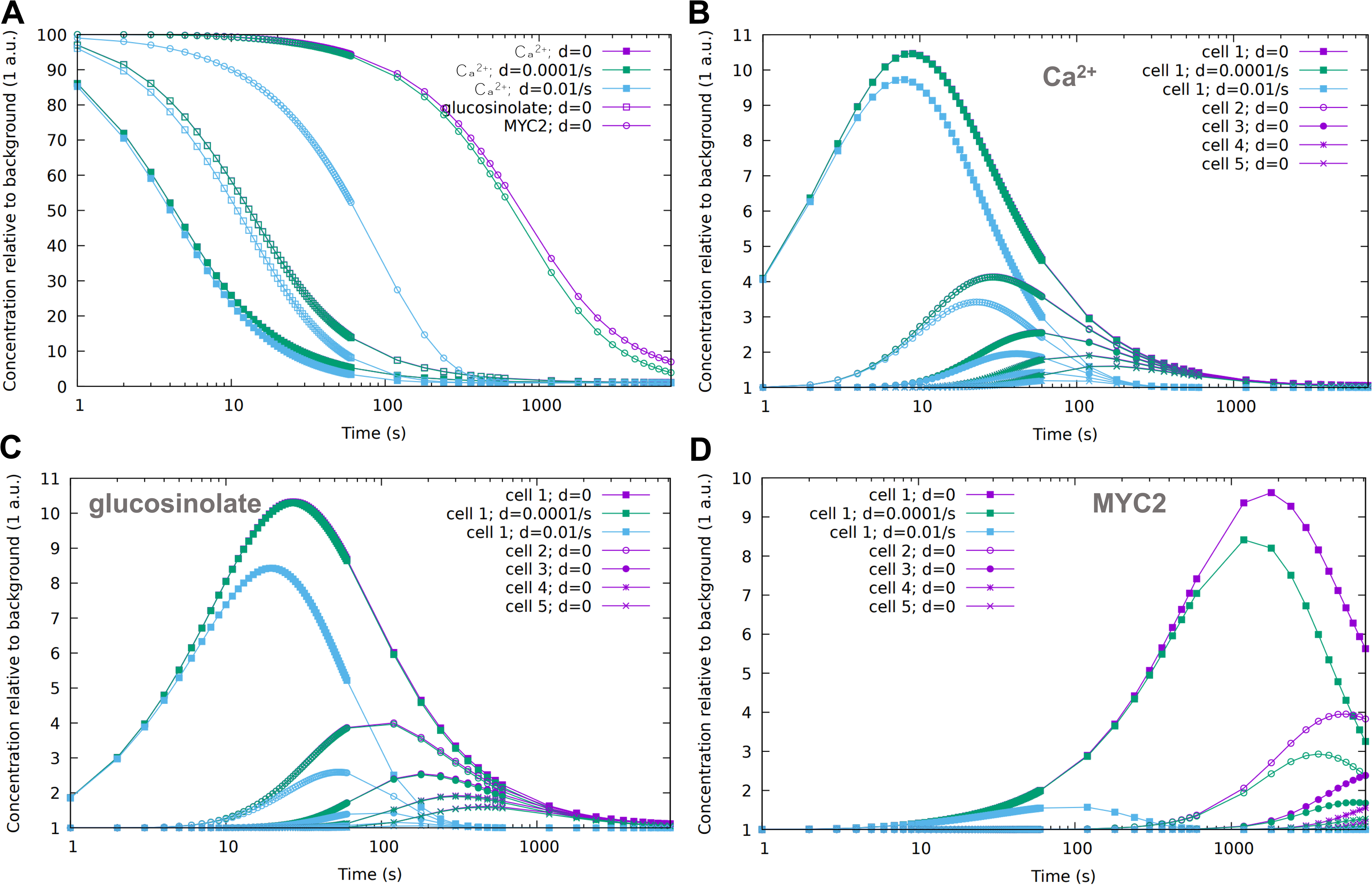
Impact of plasmodesmata permeability and turnover on the distribution range of defense compounds. Concentrations of Ca^2+^ ions, the glucosinolate 4-methylsulfinylpropyl and MYC2 were obtained through diffusion simulation for the stylized leaf described in Fig. 4. **A** Time profile of source cell concentration after an instantaneous increase (pulse) of the defense compound concentration to 100 times their respective background levels at T = 0 s. **B-D** Time profiles of Ca^2+^ ions (**B**), glucosinolate (**C**) and MYC2 (**D**) concentrations in neighbor cells (see Fig. 4C inset). Defense compound degradation rates in all cells are integrated as d = 0 (no degradation), d = 0.0001 s^-1^ (half life of almost 2 h) or d = 0.01 s^-1^ (half life of 69s). The turnover rate could also refer to sequestration. Data for continuous production of the defense compounds in the source cell is presented in Fig. S2.

**Fig. 6.**
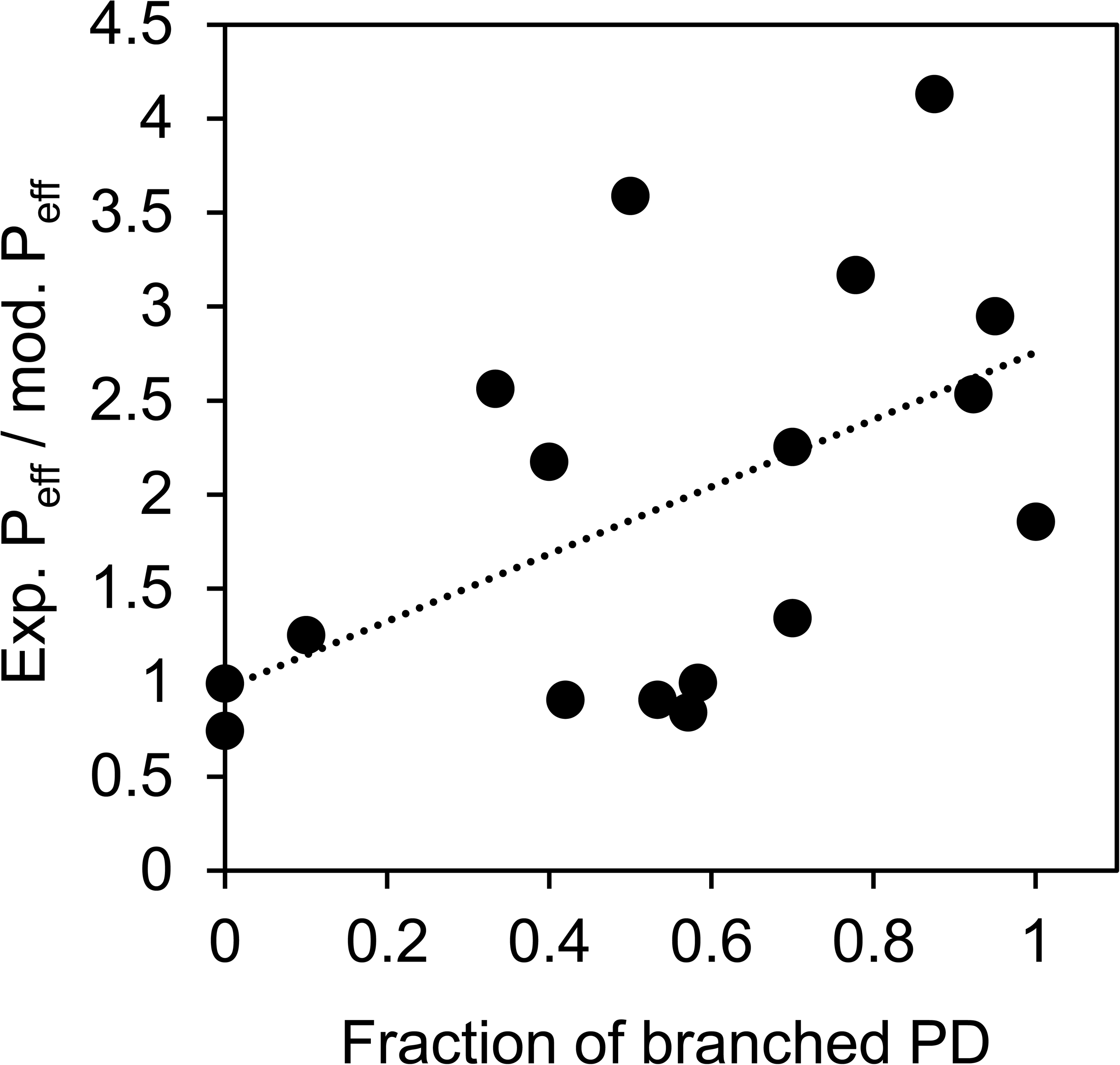
Branched plasmodesmata (PD) and the relationship of experimentally determined and modelled interface permeabilities. The ratio of experimental and modelled P_eff_ (data from Table 4) is plotted against the fraction of branched plasmodesmata. Linear regression line indicates significant correlation (R = 0.53; p = 0.019, n = 19).

We chose three molecules that are locally produced in response to herbivory attack: Ca^2+^, the glucosinolate 4-methylsulfinylpropyl (referred to as glucosinolate) and the transcription factor MYC2 (Halkier and Gershenzon 2006; Clark et al. 2016; Nguyen et al. 2018). These represent molecules of very different sizes. In hydrated state, Ca^2+^ ions have a diameter of 0.45 nm, glucosinolate of 1.2 nm and MYC2 about 7 nm. We first simulated the diffusion of the three compounds from a single source cell at the edge of the leaf without any degradation or sequestration of the molecule (Fig. 4D). These simulations show the theoretical maximum range of the signal at different time points. All substance was instantaneously released in the source cell at t=0 with a concentration of 100 times the background concentration of 1 arbitrary unit (a.u.) of all other cells. Relative concentrations were calculated in all leaf cells at various times. Concentration profiles over the whole leaf at timepoints of 1 min, 10 min and 1h are visualized in Fig. 4E-G. As expected, the small Ca^2+^ ions diffused farthest, with some ions (i.e., the 0.1% concentration increase contour) reaching the other leaf half within 1 h (Fig. 4E). Some glucosinolate molecules reached the midrib in the same time (Fig. 4F). The MYC2 proteins only diffuse across a few cell-cell-interfaces (Fig. 4G). Whereas the Ca^2+^ concentration profile became relatively flat, the bigger molecules maintained steeper concentration gradients (Fig. 4).

To further quantify the diffusion-driven spread of the three compounds, we extracted how the concentrations changed over time (Fig. 5) in a line of six cells (Fig. 4D). The plots enable an evaluation of when meaningful concentrations are reached in the cells neighboring the source cell. Moreover, we take defense compound degradation into account by providing concentration dynamics with degradation rates of 0.01 s^-1^ and 0.0001 s^-1^, equivalent to molecule half-lives of 69 s and almost 2 h, respectively. In case d > 0, all cells produce the substance at a low rate, to maintain the equilibrium background concentration of 1 a.u.. Especially in case of Ca^2+^ this degradation factor would also cover defense compound sequestration in the ER and vacuole.

Fig. 5 shows time profiles after a pulse of that increased the respective defense compound to 100 times its background level in the source cell. The plots show, for example, that the Ca^2+^ concentration in the third neighbor cell reaches double the background concentration after about 25 s after a pulse in the source cell (Fig. 5B). The glucosinolate would reach concentration doubling in the third neighbor cell after about 120 sec (Fig. 5C). In contrast, a concentration doubling in the third neighbor cell would take more than one hour for MYC2 and would only be reached at all in the absence of any degradation (Fig. 5D). Degradation, at the rates simulated here, has a relatively minor influence on the diffusion of Ca^2+^, but major effects for glucosinolate and MYC2 distributions (Fig. 5B,C).

As it may take time to build up a high concentration of a signaling molecule in the source cell (see Table 6), we also tested an alternative scenario in which the source cell starts producing signal at T = 0, which could then immediately start spreading. Plots in which defense compounds are constantly produced in the source cell are provided in Fig. S2. The curves for d = 0.01 s^-1^ show that in this case, after 2-3 half-lives of the signal, the concentrations reach a plateau. This saturation means that the degradation of the signal determines how far it can reach, and, moreover, the shorter the half-life of the signal, the faster its steady state distribution will be reached.

**Table 6.** Information on defense compound size, potential concentrations and potential time of production during defense response. Hydrodynamic radii were used to calculate cytosolic diffusion coefficients according to the Stokes-Einstein-equation relative to Ca^2+^, for which an experimentally determined value was available (Donahue and Abercrombie 1987). Values for defense compound concentrations after induction and source activity periods are estimates that were synthesized from data in multiple studies. The concentration values are estimated to vary by more than an order of magnitude, considering their dependence on species, leaf status, the kind of herbivore and many additional factors. Source activity periods can be similarly expected to vary strongly, potentially by 50 to 100 %.

| Compound | Hydodynamic diameter (nm) | Cytosolic diffusion coefficient ($\text{cm}^2/\text{s}$) | Source cell concentration | Other cell concentration | Source activity period | Reference |
| --- | --- | --- | --- | --- | --- | --- |
| $\text{Ca}^{2+}$ | 0.45 | $5.3 \times 10^{-6}$ | 2 mM | 0.1 mM | 100 sec | Knight et al. 1997; Kiegle et al. 2001; Nguyen et al. 2018 |
| GLS | 1.15 |  | 1 mM | 10 nM | 6 h | Mewis et al. 2005; Halkier and Gershenzon 2006; |
| MYC2 | 7 |  | 600 nM | 6 nM | 90 min | Clark et al. 2016; Zhang et al. 2019; Heinemann et al. 2020 |

Snapshots comparing pulse and production scenarios are shown in figures S3-S5. The snapshots show that the degradation of the signal has a big impact on its range (comparing production for low and high d), and that with high degradation, the signal vanishes quickly after the source activity stops (pulse scenario, d = 0.01 s^-1^: simulated leaves at t > 10 min are almost fully at background concentration). With low degradation, on the other hand, continuous production of low mobility compounds like MYC2 can result in very high concentrations in the source cell.

The simulations can indicate defense compound distributions resulting from intercellular diffusion in the model leaf. Adjustment of the model parameters could enable similar predictions for leaves of different sizes and shapes, as well as other organs.

## Discussion

The regulation of PDs is complex and important for a wide range of plant developmental processes and stress responses (Tee and Faulkner 2024). At the same time, PDs are hard to study and a lack of experimental avenues continues to constrain further progress (Peters et al. 2021). Our results help to consolidate the methodological toolbox for the analysis of PD function. As described in the introduction, quantification of PD-mediated interface permeability with the help of fluorescent tracers and calculations in PDinsight using TEM-derived data both carry methodological uncertainties that could increase variation or skew results. Here the FLIP values or P_eff_ values derived from photoactivation experiments correlated with the model P_eff_-values based on TEM data. This result indicates that both approaches are suitable for the quantification of interface permeability.

One remaining issue is that experimental P_eff_-values were on average about two times higher than model P_eff_-values. Values are close enough to confirm the general range of interface permeabilities, especially as it agrees with previous predictions derived from slightly different approaches (Rutschow et al. 2011). Nevertheless, it is worth exploring potential causes of the discrepancy.

The photoactivation approach has a relatively high experiment-to-experiment variation of 15 to 30% of mean values across plants grown under the same conditions and of similar developmental stage. Checking the distributions for single interfaces indicated that about 35% of P_eff_ distributions were skewed towards low values, while others were normally distributed. However, using median values instead of mean values did not improve the correlation with modelled P_eff_ and lowered the ratio of experimental and theoretical P_eff_ only slightly. A more likely source for systematic errors is the image analysis procedure. As discussed previously (Liesche and Schulz 2012; Gao et al. 2020), flux values are derived from fluorescence intensities in the cells neighboring the photoactivated cell, which means that they can be influenced by cell sizes and coupling to the next neighbor cells. Principally, tracer flux to secondary and tertiary neighbor cells, and the neighbor cells in the mesophyll layer below the epidermis, could lead to underestimation of flux values. However, analysis that included these neighbors indicated that their effect is reasonably well reflected in the current analysis procedure (Gao et al. 2020). An adaptation of methods for high-speed, high-resolution three-dimensional tracer imaging (Zanacchi et al. 2013) to leaves would be needed to clarify this issue.

Above considerations indicate that the modelled P_eff_ potentially underestimates the actual P_eff_ to a certain degree. The TEM image analysis itself is unlikely to introduce any systematic error. Our conventional sample preparation yields similar images and results compared to the potentially less invasive preparation using cryofixation and freeze substitution (Nicolas et al. 2017; Yan et al. 2019). Moreover, we found the variations in desmotubule diameter to be too small to have a major impact on calculations. Low variation in desmotubule diameter was expected since PD permeabilities in the *Pisum sativum* root cortex cells exposed to various osmotic treatments were previously found to correlate with neck diameter, while desmotubule diameters remained constant (Schulz 1995). One potential source of uncertainty is PD shape that can have significant impact on permeability (Ostermeyer et al. 2022). However, the appearance of the PDs analyzed here was relatively uniform. Another factor could be PD branching. There is not sufficient data available to relate small molecule diffusion rates to branching anatomies and, accordingly, it could not be integrated into the theoretical description of PD diffusion (Deinum et al. 2019). Our data shows that the ratio of experimental P_eff_ to modelled P_eff_, as indicator of the discrepancy between the two approaches, correlates with the fraction of branched PD at a given interface (Fig. 7). This means that branching could be an important factor for interface permeability. If branching increased permeability compared to unbranched PD of similar dimensions, its inclusion into theoretical models would help to unite the two approaches. Ideally, correlative electron and live-cell microscopy of interfaces with various fractions of branched PD would be used to map PD structure to diffusion rates.

A process in which PD undoubtedly play a major role is the plant’s response to biotic stress (Lim et al. 2016). Interface permeability calculation with PDinsight can facilitate a quantitative view, because it can provide results for molecules of various sizes (Deinum et al. 2019). We chose three molecules that are locally produced in response to herbivore attack to illustrate this capacity. These molecules, Ca^2+^ ions, glucosinolate and the transcription factor MYC2, play important roles in organ-level signaling (Table 6), meaning that quantification of their movement is important for understanding the molecular mechanism of the response. Our simulations predict the spread of these molecules at a physiological range of concentrations, source activity periods and turnover rates (Table 6). Thereby, it can provide a quantitative basis for the integration of intercellular diffusion rates with trans-membrane transport, apoplasmic transport and other factors that influence intercellular movement. For example, our inclusion of molecule degradation in the simulations indicates that degradation is more limiting on the range of small defense compounds than PD permeability, in line with 1D calculations by Deinum (2013). Such integration will be essential for filling our knowledge gap on the relative contributions of PD and trans-membrane transport that currently limit our understanding of the movement of auxin in the leaf hyponasty response (Li et al. 2023) or the movement of metabolites between Kranz mesophyll cells and bundle sheath cells in C4-type photosynthesis (Bilska and Sowinski 2010).

## Material and Methods

### Plant material

Arabidopsis plants (ecotype Col-0) were grown in growth chambers at 16 h light/8 h dark periods, with light levels of 200 μmol photons m^-2^ s^-1^ and constant temperature of 21 °C. Arabidopsis t-DNA insertion mutant *gsl8* (NCBI stock number N665827) was grown under the same condition as wild-type plants. *N. tabacum* plants were grown in growth chambers at 350 μmol photons m^-2^s^-1^ irradiation, 16 h light/8 h dark periods and temperatures of 25 °C. Leaves of *N. tabacum* were sampled 28 days after germination. *Pisum sativum* L. cv. “Kleine Rheinländerin” seeds were soaked in aerated tap water for one day before cultivation on wet vermiculite. Plants were grown in growth chambers with 18 h illumination at 200 μmol photons m^-2^s^-1^ and 6 h dark at 24 °C. *Populus x canescens* leaves were sampled from a 10 m high tree (about 12 years old) at the Botanical Garden of Copenhagen University (Latitude 55.67, Longitude 12.46) on sunny spring (May) days with temperatures between 16°C and 22°C.

### Transmission electron microscopy

The results of TEM analysis regarding PD structure and density were extracted from Zhu et al. 1998, Liesche et al. (2019) and Gao et al. (2020) for most interfaces. Only data for the Arabidopsis epidermis cells in young and transition stage leaves, the mesophyll cells in mature Arabidopsis leaves and the root cortex cells of *P. sativum* were generated as part of this study. For the TEM analysis of Arabidopsis, the protocol of Gao et al. (2020) was precisely followed. This means that sections of 2-3 mm width were cut from leaves and fixed in 2.5% glutaraldehyde in 0.1M phosphate buffered saline (PBS, pH 7.2) for at least 12 hours at 4°C. Samples were washed four times in 0.1M PBS (pH 7.2) for 15 minutes before incubation in 1% osmic acid for 4 hours at 4°C. Samples were then washed in 0.1M PBS (pH 7.2) before dehydration in an acetone series. Samples were embedded in LR White resin (Sigma-Aldrich, St. Louis, MO, USA), followed by trimming (EM TRIM2, Leica Microsystems, Wetzlar, Germany) and sectioning on an ultramicrotome (UC7, Leica Microsystems) with section thickness of 70 nm. Sections were transferred to copper grids and consecutively stained in 2% uranyl acetate for 20 minutes and lead citrate for 15 minutes. TEM imaging was performed on a Fei TECNAI G2 SPIRIT BIO (Hillsboro, OR, USA).

*P. sativum* seedling preparation and imaging followed the protocol of Schulz (1995). Root tips were cut and immediately immersed in a Karnovsky fixative at pH 7.3. After rinsing with 100 mM Na-cacodylate buffer, samples were post-fixed in 1% OsO_4_, dehydrated through an acetone series and embedded in Spurr’s resin, followed by polymerization at 60°C for 36 h. Ultrathin sections of about 70 nm thickness were mounted on formvar-coated single slot grids and post-stained with 1% methanolic uranyI acetate for 15 min. These were rinsed in a methanol/H_2_0 series, before incubation in aqueous lead citrate for 5 min. Images were collected on a Talos L120C TEM (Thermo Scientific).

PD density D was calculated from electron micrographs as recommended by Robards (1975) with the formula D = count per µm wall length / (T + 1.5r) with section thickness T = 70 nm and average PD radius r of the interface in question. Any significant portion of a PD visible in a micrograph (typically at least ¼ PD) was included in the quantitative analysis.

### Live-cell microscopy

FLIP values were extracted from Liesche et al. (2019). Results of photoactivation experiments were obtained from Liesche and Schulz (2012) for *Cucurbita maxima* leaf epidermis cells and from Gao et al. (2020) for Arabidopsis epidermis cells in mature leaves. As part of this study, photoactivation experiments were conducted on Arabidopsis epidermis cells in leaves at young and transition stages. The general parameter for categorization of developmental stages was leaf size. Young leaves are below 2 mm across, transition stage leaves 3-4 mm and mature leaves have a width of more than 5 mm. Furthermore, experiments were carried out on leaf epidermis cells of *Nicotiana tabacum* and *Populus x canescens* and various cell-cell interfaces in Arabidopsis roots.

Our experimental procedure is described in detail Gao et al. (2020). That publication also contains results of control experiments that were conducted to support the method. The procedure of Gao et al. (2020) was used for the current study with some changes. Instead of caged fluorescein, we used Cage500 (Abberior, Göttingen, Germany) as photoactivatable tracer. Cage500 has very similar properties to caged fluorescein, including similar size, but can be activated by the common 405 nm laser instead of the non-standard 355 nm laser (Belov et al. 2014). The tracer was loaded into Arabidopsis leaf tissue by applying a drop of 1 mM tracer solution to a 1 mm incision made using a razor blade at the edge of a leaf. *N. tabacum* and *P. canescens* leaves were prepared similarly, only that the incision was about 0.5 to 1 cm on their much bigger leaves. Arabidopsis roots were submerged in tracer solution. All samples were imaged after incubation for 20 min at room temperature.

Confocal laser scanning microscopes, Leica Microsystems (Wetzlar, Germany) model SP2 for roots and model SP8 for leaves, were used. The two instruments were equipped with similar 405 nm and 488 nm lasers, similar non-resonant scanner and similar photomultiplier tube (PMT) detectors, yielding virtually identical results. It should be noted that, as long as photoactivation times are in the range of 1 to 20 sec, interface permeabilities calculated from photoactivation experiments are hardware-independent (Gao et al. 2020). A 40x water-dipping objective (NA = 0.95) was used for all experiments. Fluorescence of the activated tracer was monitored before, during and after photoactivation by excitation at 488 nm (20% laser power of 100 mW laser) and emission detection between 495 and 550 nm. A target cell was selected and outlined using the corresponding bright field image. The 405 nm diode laser at maximal output power (40 mW) was used for photoactivation of the target cell. Gain of the photomultiplier tube detector was set to somewhat below signal saturation, typically at 700 to 800 V, and two-times line averaging was activated. The duration of the photoactivation phase varied between 2 and 10 sec according to sample type.

Image processing, analysis and calculation of interface permeability followed exactly the protocol described by Gao et al. (2020). Three regions of interest (ROIs) were drawn to determine the flux of the activated dye and to quantify the concentration potential for the interfaces between the target cell and each neighboring cell, using ImageJ (Schindelin et al. 2012): For flux measurement, a ROI corresponding to the respective neighbor cells was drawn and values for total fluorescence (integrated density) were extracted. For dye concentration measurements, a ROI of about 5 µm width was drawn on each side of the respective interface and values for relative fluorescence (integrated density divided by area) were extracted. In addition, the length of the interface between target and neighbor cells was measured. Using these parameters, the flux and permeability between the target cell and each neighboring cell was determined according to the following procedure. The background-corrected neighbor cell fluorescence after photoactivation was normalized to the photoactivation time to yield values for flux (in mol µm^-2^ s^-1^). To determine the concentration potential, the value of relative fluorescence in the ROI on the neighbor-cell side of the respective interface is subtracted from the relative fluorescence on the target-cell side. Assuming a linear increase between pre- and post-photoactivation states, the post-activation concentration potential was divided by two to represent the average concentration potential (see discussion in Liesche and Schulz 2013). The interface permeability P (in µm s^-1^) was then calculated by dividing the flux F by the concentration potential across the interface: P = F / Δc.

### Calculation of effective wall permeability

Effective wall permeabilities were calculated using PDinsight version 2.0 (Deinum et al. 2019; Deinum 2022) using the “singleInterface” mode for obtaining center values and “bootstrapInterface” for confidence intervals. We used 10000 bootstrap samples to obtain 95% confidence intervals. Pdinsight is available from https://github.com/eedeinum/PDinsight.

Neck radius (Rn, Rn2), neck length (Ln, Ln2), PD length (Lpd) and maximum radius (Rc) of individual PDs were determined from TEM images. Potential outliers were identified in a 2-stage procedure. First, all PDs with a parameter value more than 1.5 times the inter-quartile range (IQR) outside the IQR of the respective parameter of the respective data set were marked as potential outlier. Of those, the PDs that appeared outside the range of all PDs on a 2-component PCA plot were marked as likely outlier. For all likely outliers (and all other points that stood out from visual inspection of the raw numbers), we revisited the original TEM images to check for measurement errors. All (possibly corrected) points were included in the analysis, because of the relatively small sample sizes.

Density was estimated as a single number per data set (species/interface combination; see above). We assumed a hexagonal grid of PD positions and a uniform desmotubule radius of Rdt = 8.25 nm. By default, we assumed no clustering of PDs (nPit = 1), but also tested the effect of mild to moderate clustering with a cluster size of nPit = 3 and nPit = 7 and within cluster PD-PD distance dPit = 100 nm.

#### Whole-mount staining and simulation of defense compound diffusion

Arabidopsis leaves selected for whole-mount staining were incubated in fixative solution (50% methanol, 10% acetic acid, 40% water) for 12 hours at 4 °C. The leaves were transferred to 80% ethanol for 1 minute at 80 °C and moved back to fresh fixative solution for 1 hour. Leaves were washed in water and incubation in 1% periodic acid for 40 minutes at room temperature. After washing in water, leaves were incubated in Schiff’s reagent (Carl Roth GmbH. Karlsruhe, Germany) with 100 μg/mL propidium iodide (Carl Roth GmbH) for 2 hours. Samples were transferred to microscope slides and covered with chloral hydrate solution overnight. Chloral hydrate solution was prepared by mixing 4 g chloral hydrate (Carl Roth GmbH), 1 mL glycerol and 2 mL H_2_O. The chloral hydrate solution was removed and Hoyer’s solution was dropped on the sample, then covered with a microscope cover slip. Hoyer’s solution was prepared by mixing 30 g gum arabic, 200 g chloral hydrate, 20 mL glycerol and 50 mL H_2_O. Samples were imaged on a Leica TCS SP8 confocal microscope (Leica Microsystems, Wetzlar, Germany) using the tile scan function. Images were combined and analyzed in ImageJ.

A leaf epidermis template was made to represent the whole mount-imaged Arabidopsis leaf. Cells were placed in a hexagonal alignment and had an epidermal surface area of 50.6×50.6 µm^2^, represented as 11×11 pixel squares. Cells were separated by a single row of wall pixels. Epidermis cells above the mid vein consisted of multiple joint squares in a row (without intervening walls), with total lengths representative of experimental images, with aspect ratios increasing towards the leaf base/petiole.

Permeabilities of different interfaces were multiplied by different factors to compensate for 1) the fact that the transverse and longitudinal interfaces were not equally long in number of pixels between cells (q_t_ = 1.1), 2) the difference between the straight interfaces between simulation cells and the lobed interfaces of the puzzle piece-shaped epidermal cells (q_w_ = 1.8).

Intracellular diffusion coefficients (D) were based on a value of 530 µm^2^/s for Ca^2+^ (Donahue and Abercrombie 1987) and adjusted to other substances using the Stokes-Einstein relation, which predicts that D ∼ 1/r, with r the particle radius.

Base effective wall permeability for symplasmic transport (P_base_) was computed using PDinsight using calculated particle radii (Table 6) and diffusion coefficients (see above) and the measured PD dimensions from the Arabidopis EC-EC interface and a presumed desmotubule radius Rdt = 8.25 nm. This resulted in the following values of P_base_ 2.202 (Ca^2+^), 0.686 (glucosinolate) and 0.049 (MYC2). The final P_eff_ per interface was: P_base_ × q_t_ (if transverse) × q_w_ (if puzzle-puzzle interface). The source of the signal always was a single cell at the bottom edge of the simulation template (see Fig. 4D). In the pulse scenario, the starting concentration was 100 a.u. in the source cell and equilibrium background level (1 a.u.) in all other cells. Degradation (d) rate was set at one of three levels: 0 (none), 0.0001 /s (low, resulting in a half-life of roughly 2 hours), 0.01 /s (high, resulting in a half-life of 69 s). A low level of production of d × a.u. occurred everywhere to ensure the equilibrium background of 1 a.u. was maintained (rates of 0, 0.0001 a.u./s, 0.01 a.u./s, respectively).

In the (constant) production scenario, the starting concentration was set to the equilibrium background concentration of 1 a.u. everywhere. Production rate in the source cell was 1 a.u./s for the full duration of the simulation and d × a.u. in all other cells (see above).

All simulations were performed using the same simulation software as (Deinum et al 2016), using the adaptations for symplasmic transport between cells as described in (Deinum 2013). The simulations consider both intracellular diffusion and passive transport through the walls of neighbouring cells via the PDs. The software uses a first order Alternating Direction Implicit (ADI) numerical solver (Peaceman and Rachford 1955) with an integration time step of 0.1 s (simulation time up to 60 s), 0.5 s (simulation time up to 9 min) or 1 s (simulation time up to 2 h). Whole leaf concentration profiles were measured every 1s, 1 min, or 9 min for simulation times up to 1 min, 10 min, 2 h, respectively. All simulations were started from t = 0 s. Where applicable, the simulations with the shortest integration time step were used for time traces and concentration profiles.

### Quantification and statistical analysis

The statistical analysis related to calculating PD permeability were performed using R, gnuplot, Microsoft Excel software or Python code (PDinsight). All experiments to quantify interface permeability had at least six biological replicates, with samples from at least three plants. Welch’s t test was performed at a significance level of p < 0.05.

### Data and code availability

The published article includes all datasets generated or analyzed during this study, including the raw data on PD structure and abundance (Table S2). Relevant code is available on GitHub under https://github.com/eedeinum/PDinsight.

## Acknowledgements

We thank Ms Wang Zhen, Mr Huang KeRang, Ms Chen Lei and Ms Liu XiaoRui (Life Science Research Core Services, Northwest A&F University, Yangling, China) for their assistance in the use of transmission electron microscopy at Northwest A&F University. Publication costs were covered by the University of Graz.

## Author contributions

This study was conceptualized by Johannes Liesche and Eva Deinum. Experiments were planned by Johannes Liesche, Helle Martens, Alexander Schulz and Eva Deinum and carried out by Jiazhou Li, Helle Martens, Chen Gao and Alexander Schulz. Data analysis was carried out by all authors. Johannes Liesche and Eva Deinum drafted the manuscript, which all authors helped to finalize.

